# Late Quaternary millennial-scale stability of the feeding strategy of the gray brocket deer [*Subulo gouazoubira* (G. Fischer, 1814); Mammalia] in Southeastern Brazil

**DOI:** 10.1101/2024.11.08.621301

**Authors:** William J Pestle, Alex Hubbe, Mark Hubbe, Avaaz Amirali, Eliane Nunes Chim, Rodrigo Elias Oliveira, Eluzai Dinai Pinto Sandoval, Lorena Becerra-Valdivia, José Maurício Barbanti Duarte, Walter A Neves

## Abstract

There is only limited evidence (contemporary or archaeological/paleontological) regarding the feeding ecology of the gray brocket deer (*Subulo gouazoubira*) in Brazil’s Cerrado biome. Using stable carbon (δ13C) and oxygen (δ18O) isotope analyses of dental enamel from individuals recovered from Cuvieri Cave, Minas Gerais, we explore dietary and environmental changes between the Late Pleistocene and Holocene epochs for *S. gouazoubira* living in this biome. δ13C data suggest no significant shift in feeding strategy between the Late Pleistocene and Holocene, with both sets of individuals displaying carbon isotope values consistent with browsing in open environments rather than closed canopy forests. A shift in δ18O values, on the other hand, indicates climatic changes in the area of Cuvieri Cave from the late Pleistocene to Holocene epoch. This study highlights the stability in the feeding behavior of the gray brocket deer despite biotic and abiotic changes between the epochs.

## Introduction

The gray brocket deer [*Subulo gouazoubira* (G. Fischer, 1814); Mammalia; Bernegossi et al., 2023] is a small- to medium-sized deer (weight 11–25 kg) distributed primarily in tropical, subtropical, and semiarid regions south and east of the Amazon forest (Figure 1). It inhabits moderately humid to dry areas with some degree of tree cover, avoiding dense rainforests but frequently occupying mosaics of open savannas, dry forests, and shrublands (Black-Décima & Vogliotti, 2016). The species is particularly common in ecotones between open and wooded formations, such as the Cerrado *sensu lato*, Chaco woodlands, and Caatinga scrublands, where it forages on a diverse assemblage of herbaceous and shrubby vegetation, complementing its diet with fungi, feces, invertebrates, bones, and soil (see Black-Décima & Vogliotti, 2016; Black-Décima et al., 2010, and references therein). Its diet varies throughout the year and across geographic locations according to food availability, and feeding activity often occurs in open patches adjacent to cover, reflecting a browsing–mixed feeding strategy typical of cervids adapted to heterogeneous landscapes (Black-Décima & Vogliotti, 2016; Gordon & Prins, 2019; Grotta-Neto et al., 2024).

**Figure 1.**
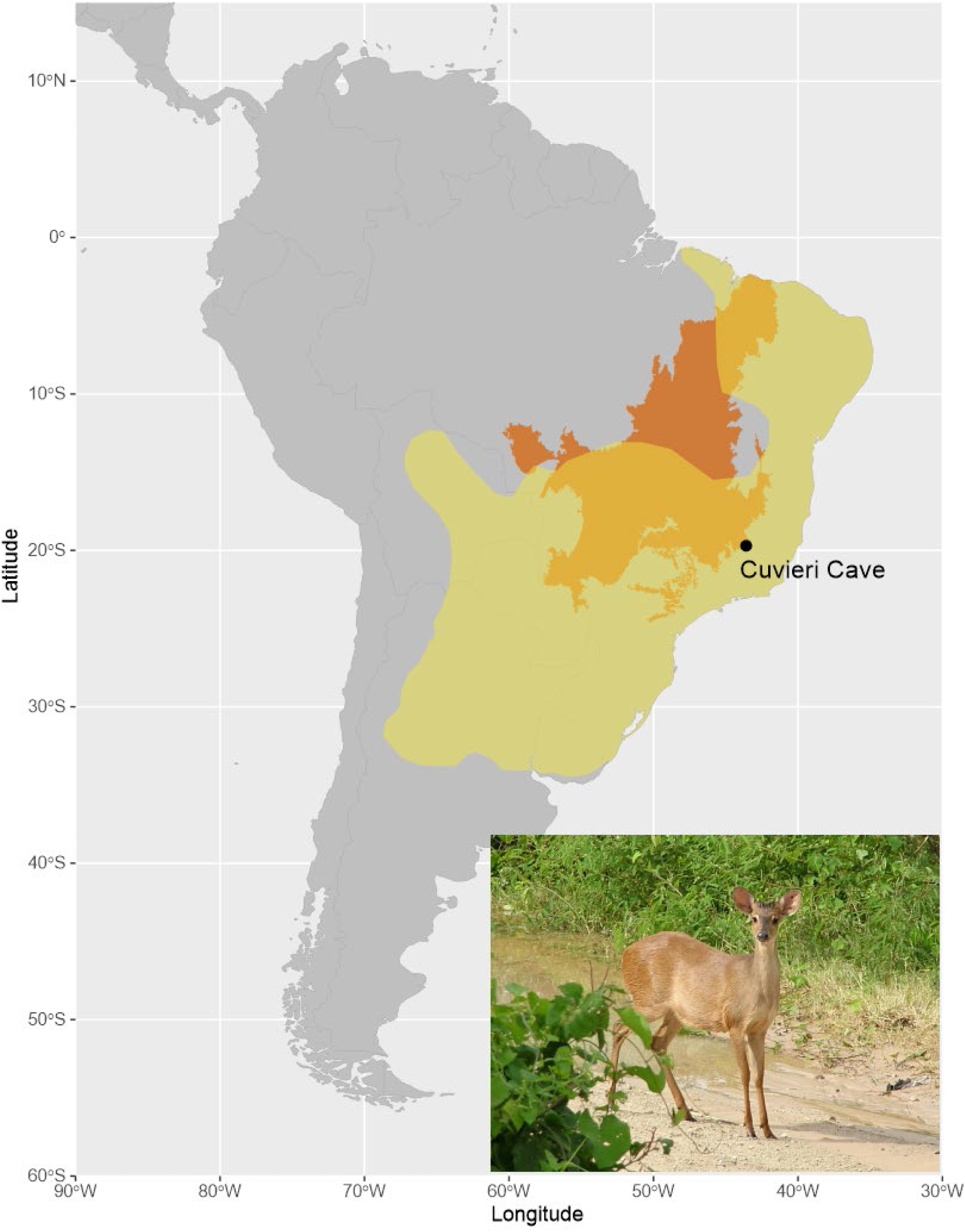
The current distribution range of the gray brocket deer (*Subulo gouazoubira*; yellow area) and Cerrado domain (orange area). The lower right picture depicts an adult male gray brocket deer. The locality of Cuvieri Cave is shown on the map. Gray brocket deer distribution based on IUCN (2008). Cerrado distribution based on IBGE (2015). Photo by: José Maurício Barbanti Duarte.

In the Brazilian savanna (i.e., Cerrado domain; Klink & Machado, 2005; Ribeiro & Walter, 2008), where the gray brocket deer is considered an abundant species (Marinho-Filho et al., 2002), there are no studies focused on its current or past diet. Given the species’ dietary flexibility, it is plausible that its diet has varied during the end of the Quaternary for two main reasons. First, there is evidence of vegetation and climate changes spanning at least the last 30,000 years (Azevedo et al., 2021; Bissa & Toledo, 2015; Cheliz et al., 2021; Novello et al., 2019; Raczka et al., 2017; Turcq et al., 1997). Second, several species went extinct during the late Quaternary (Cartelle, 1999), which may have altered direct and indirect ecological interactions. At a minimum, a horse (*Equus neogeus* Lund, 1840), two ground sloths (*Catonyx cuvieri* Lund, 1839, and *Valgipes bucklandi* Lund, 1844), and the saber-toothed cat (*Smilodon* populator Lund, 1839) became extinct after ∼20,000 cal yr BP (calibrated years before present, with ‘present’ defined in ^14^C dating as 1950 in the common era), with the last three species having final appearance dates younger than ∼13,500 cal yr BP (Hubbe et al., 2013).

For nearly fifty years, the analysis of the stable isotopes of light elements (particularly carbon and oxygen) has been an important means for reconstructing the diet and ecology of ancient animals (see reviews in Clementz, 2012; Koch, 2007). While made more difficult by the sometimes-complex physiologies of the taxa under study, preservation, and diagenetic effects, stable isotope analysis can provide crucial insights into animal behavior spanning archaeological and geological time. In particular, owing to its composition and structure (Lee-Thorp, 1989; LeGeros, 1991), the carbon and oxygen isotope makeup of the carbonate fraction of dental enamel bioapatite has been demonstrated to preserve biogenic signals of feeding ecology and paleotemperature/paleohydrology stretching back hundreds of millions of years (Koch 2007: Table 5.2).

In this study, we employed stable isotope analysis (δ^13^C and δ^18^O) on dental enamel bioapatite to explore potential variations in the feeding strategy of gray brocket deer and their environment during the late Quaternary within the Cerrado, using samples obtained from Late Pleistocene and Holocene deposits at Cuvieri Cave, Lagoa Santa, Minas Gerais, Brazil (Figure 1; Hubbe et al. 2011; Haddad-Martim et al. 2017). Specifically, we sought to characterize the feeding ecology, habitat type, and paleoclimatic context of this taxon, comparing samples drawn from Pleistocene and Holocene levels in the cave deposits. Given the small sample sizes involved, we recognized from the outset that these results are preliminary and contingent and should be verified by further research. However, given the dearth of information about the species diet in the past, our results are relevant to contextualize a species that has been shown great resilience in their habitats (Hubbe et al., 2025).

## Materials and Methods

All the specimens analyzed are stored in the collection of the Laboratory of Human Evolutionary Studies, Institute of Biosciences, University of São Paulo, Brazil. Essential information regarding the analyzed samples from the Cuvieri Cave fossil site is presented here. For further information on the Cuvieri Cave fossil site, including its stratigraphy and chronology, please refer to Hubbe et al. (2011, 2025) and Haddad-Martim et al. (2017).

Four lower right third molars were sampled from late Pleistocene facies (>30,000 yr) and 25 lower left third molars were sampled from Holocene facies (<10,755 cal BP) of Cuvieri Cave (Table 1). The discrepancy in sample sizes between the Late Pleistocene and the Holocene reflects the abundance of teeth available and is not the result of an intentional sampling strategy.

**Table 1.** Anatomical, chronostratigraphic, and isotopic data for dental enamel samples from Cuvieri Cave’s Holocene and Pleistocene deposits. The “depth” column refers to the vertical distance between the location where the sample was found during excavation and the adopted zero reference point in the excavated areas. Pleistocene samples (indicated with *) lack sufficient chronological resolution to model dates and are thus conservatively treated to be older than 30,000 cal BP following Haddad-Martim et al. (2017).

| Excavation # | Anatomical part | Depth (cm) | Epoch | Modeled age (95% CI, cal BP) | Modeled age (median, cal BP) | Apatite yield | $\delta^{13}\text{CPDB}$ (‰) | Diet maximum $\delta^{13}\text{C}$ (fractionation -12‰) | Diet minimum $\delta^{13}\text{C}$ (fractionation -14.6‰) | $\delta^{18}\text{OPDB}$ (‰) | $\delta^{18}\text{OSMOW}$ (‰) Coplen 1983 | $\delta^{18}\text{Odw}$ (‰) Stevens, et al. 2011 |
| --- | --- | --- | --- | --- | --- | --- | --- | --- | --- | --- | --- | --- |
| CVL3B 352 | Hemimandible | -112.5 | Pleistocene | n/a | 30000* | 71.5% | -10 | -22 | -24.6 | -4.7 | 26 | -7.6 |
| CVL3B P2895 | Hemimandible | -130<sup>a</sup> | Pleistocene | n/a | 30000* | 77.9% | -13.4 | -25.4 | -28 | -5.5 | 25.3 | -8.4 |
| CVL3B P3961 | isolated tooth | -168<sup>a</sup> | Pleistocene | n/a | 30000* | 74.8% | -10.3 | -22.3 | -24.9 | -5.2 | 25.5 | -8.1 |
| CVL3B P6143 | isolated tooth | -257<sup>a</sup> | Pleistocene | n/a | 30000* | 84.1% | -10 | -22 | -24.6 | -4.7 | 26 | -7.6 |
| CVL2 687 | Hemimandible | -66 | Holocene | 2943-1370 | 2295 | 86.6% | -11.3 | -23.3 | -25.9 | -1.4 | 29.4 | -4.2 |
| CVL2 2140 | Hemimandible | -70.5 | Holocene | 3862-2810 | 3342 | 69.9% | -10.4 | -22.4 | -25 | -3.2 | 27.6 | -6.0 |
| CVL2 3361 | Hemimandible | -75 | Holocene | 5875-3713 | 5098 | 62.0% | -12.6 | -24.6 | -27.2 | -3 | 27.8 | -5.8 |
| CVL2 5958 | Hemimandible | -75 | Holocene | 5875-3713 | 5098 | 66.7% | -10.6 | -22.6 | -25.2 | -2.2 | 28.7 | -5.0 |
| CVL2 2957 | Hemimandible | -75 | Holocene | 5875-3713 | 5098 | 74.1% | -12.2 | -24.2 | -26.8 | -2.7 | 28.1 | -5.5 |
| CVL2 4321 | isolated tooth | -76 | Holocene | 5943-3786 | 5767 | 74.9% | -10.9 | -22.9 | -25.5 | -2.9 | 27.9 | -5.7 |
| CVL2 6730 | Hemimandible | -80.5 | Holocene | 5944-5611 | 5835 | 63.6% | -10.5 | -22.5 | -25.1 | -2.6 | 28.3 | -5.4 |
| CVL2 6912 | Hemimandible | -88 | Holocene | 7878-5787 | 6994 | 70.4% | -11.8 | -23.8 | -26.4 | -3.4 | 27.5 | -6.2 |
| CVL2 8210 | isolated tooth | -88.5 | Holocene | 7878-5886 | 7182 | 70.8% | -10.6 | -22.6 | -25.2 | -1.7 | 29.2 | -4.5 |
| CVL2 10743 | Hemimandible | -99 | Holocene | 8251-7621 | 7827 | 71.3% | -13.7 | -25.7 | -28.3 | -3.1 | 27.8 | -5.9 |
| CVL2 10801 | Hemimandible | -99 | Holocene | 8251-7621 | 7827 | 67.7% | -10.2 | -22.2 | -24.8 | -2.7 | 28.1 | -5.5 |
| CVL2 10823 | isolated tooth | -99 | Holocene | 8251-7621 | 7827 | 68.7% | -11.3 | -23.3 | -25.9 | -3.4 | 27.4 | -6.3 |
| CVL2 10981 | Hemimandible | -100 | Holocene | 8288-7634 | 7861 | 72.3% | -10.7 | -22.7 | -25.3 | -4.4 | 26.3 | -7.3 |
| CVL2 10998 | Hemimandible | -100 | Holocene | 8288-7634 | 7861 | 66.2% | -9.8 | -21.8 | -24.4 | -3.9 | 26.9 | -6.7 |
| CVL2 12417 | Hemimandible | -102 | Holocene | 8356-7677 | 7938 | 55.8% | -8.9 | -20.9 | -23.5 | -2.1 | 28.7 | -4.9 |
| CVL2 11439 | isolated tooth | -102 | Holocene | 8356-7677 | 7938 | 79.1% | -8.6 | -20.6 | -23.2 | -4.5 | 26.3 | -7.4 |
| CVL2 12184 | isolated tooth | -104 | Holocene | 8403-7714 | 8020 | 76.9% | -8.9 | -20.9 | -23.5 | -3.5 | 27.3 | -6.4 |
| CVL2 12808 | isolated tooth | -106 | Holocene | 8450-7760 | 8104 | 74.7% | -8.4 | -20.4 | -23 | -2.3 | 28.5 | -5.1 |
| CVL2 11942 | Hemimandible | -107 | Holocene | 8470-7792 | 8146 | 74.2% | -10.9 | -22.9 | -25.5 | -2.6 | 28.3 | -5.4 |
| CVL2 12646 | isolated tooth | -107 | Holocene | 8470-7792 | 8146 | 79.9% | -9.6 | -21.6 | -24.2 | -3 | 27.8 | -5.8 |
| CVL2 12945 | isolated tooth | -108 | Holocene | 8491-7823 | 8188 | 69.6% | -11 | -23 | -25.6 | -4.9 | 25.9 | -7.8 |
| CVL2 13197 | isolated tooth | -109 | Holocene | 8509-7859 | 8231 | 79.8% | -10.1 | -22.1 | -24.7 | -2.3 | 28.6 | -5.1 |
| CVL2 12556 | Hemimandible | -110 | Holocene | 8525-7897 | 8272 | 61.8% | -10.9 | -22.9 | -25.5 | -2 | 28.9 | -4.8 |
| CVL2 12555 | isolated tooth | -110 | Holocene | 8525-7897 | 8272 | 74.9% | -10 | -22 | -24.6 | -3.8 | 27.0 | -6.6 |
| CVL2 14618 | Hemimandible | -114 | Holocene | 8586-8121 | 8411 | 73.1% | -9.7 | -21.7 | -24.3 | -3.6 | 27.2 | -6.5 |
|  |  |  |  |  | Mean | 72.2% | -10.6 | -22.6 | -25.2 | -3.3 | 27.5 | -6.1 |
|  |  |  |  |  | sd | 6.8% | 1.3 | 1.3 | 1.3 | 1.1 | 1.1 | 1.1 |
|  |  |  |  |  | Minimum | 55.8% | -13.7 | -25.7 | -28.3 | -5.5 | 25.3 | -8.4 |
|  |  |  |  |  | Maximum | 86.6% | -8.4 | -20.4 | -23.0 | -1.4 | 29.4 | -4.2 |

The teeth selected were not identified *a priori* to the species level given the difficulties in determining species based on dental morphology alone. However, we conclude that at least the majority, if not all of them belong to *S. gouazoubira* for the following reasons:

First, from a total 81 individuals identified as *Mazama/Subulo* for the Holocene and Pleistocene deposits from Cuvieri Cave [based on the minimum number of individuals (MNI); Hubbe et al., 2025], 86% (64 out of 74) of the individuals from the Holocene and 57% (4 out of 7) from the Late Pleistocene were positively identified as gray brocket deer. The remaining individuals (13) were not identified at the species level because available bones were too fragmented or polished for such identification. Thus, it is likely that most individuals not identified to the species level represent gray brocket deer, and that the sampled teeth are probably from this species.

Second, even if we assume that the individuals not identified at the species level are red brocket deer (*M. americana*), our samples from the Holocene and the Late Pleistocene likely include several gray brocket deer individuals. We consider the MNI to be a good estimator of the total amount of individuals in each time interval because: i) most specimens [i.e., a bone or tooth, or their fragment; (Lyman, 1994)] found in Cuvieri Cave deposits are the results of pitfall entrapment; ii) around 75% of the area of the Holocene and 78% of the area of the Late Pleistocene deposits were excavated; and iii) for the Late Pleistocene deposit, almost all the volume of the fossiliferous sedimentary facies were excavated, and for the Holocene deposit a volume of 3.3 m^3^ was excavated (Haddad-Martim et al., 2017; Hubbe et al., 2011). Although we did not reach the bottom of the younger deposit, vertical stratigraphic displacement is limited (Hubbe et al., 2011, 2025) and, consequently, the MNI represents most individuals from the excavated sediments. Thus, considering that the MNI is a good approximation of the sample space of deer for each epoch, it is extremely unlikely we would have sampled other species even if red brocket deer were part of the original animal assemblage. In the most conservative case, if all the unknown deer from the Holocene strata were red brocket deer (10 out of 74), the chance of sampling 19 or more gray brocket deer from this assemblage is ∼98%. Similarly, even if all unclassified specimens from the Late Pleistocene deposit (3 out of 7) were red brocket deer chance of sampling two or more gray brocket deer is ∼88%.

The sampled teeth represent individuals that were already consuming an adult diet. While direct evidence from wild populations is lacking, observations from captive animals indicate that weaning concludes between the fourth and sixth month of life (Baldini and Duarte, 2020). Tooth mineralization likely begins between 10 and 12 months of age and is generally completed by 18 to 24 months, starting at least four months after weaning and lasting approximately 6 to 14 months. A relevant caveat is that, to our knowledge, direct evidence for M3 mineralization in *S. gouazoubira* is unavailable. We therefore assumed that data from other small- to medium-sized cervids (e.g., *Odocoileus virginianus*; Severinghaus, 1949; Demarais & Strickland, 2011) provide a reasonable proxy.

### Bayesian chronological modelling

For the undated Holocene samples, the published chronological data of the respective facies at Cuvieri Cave were used to estimate sample ages through Bayesian chronological modelling. Eleven radiocarbon dates obtained from collagen (BETA-202779, -205334, -251075, -235460, - 202780, -251078, -218173, -251079, -261274, -230973, -248058; see Table 4 in Hubbe et al., 2011) were entered into a Poisson deposition model with a k parameter (k₀) of 100, an interpolation rate of 100 (i.e., an output every 1 cm), and a log₁₀(k/k₀) prior of U(−2, 2) (Ramsey, 2008; Bronk Ramsey & Lee, 2013). The latter parameter allows k/k₀ to vary by two orders of magnitude in either direction (0.01–100). The ‘General’ outlier model was also applied to all dates, with each assigned a 5% prior probability of being an outlier (Bronk Ramsey, 2009b). A radiocarbon date at 220±40 bp (BETA-205335; see Table 4 in Hubbe et al., 2011) was not included in the analysis as a clear outlier. Using the model output, the age of each sample was inferred from the depth recorded during excavation at the closest point at the respective collection level. In contrast, for the undated Pleistocene samples, the available chronological resolution is insufficient to allow age estimation. These are therefore conservatively treated to be older than 30,000 cal BP following Haddad-Martim et al. (2017). The OxCal code used for the Bayesian modeling is reported in Supplementary Information 1.

### δ^13^C and δ^18^O Isotope Analyses

Carbon and oxygen stable isotope analysis was performed on the carbonate portion of bioapatite extracted from dental enamel. Both isotope measures are well-established in archaeology, geology, and paleontology, with decades of supporting literature, and are widely used for the reconstruction of ancient animal diet, paleoecology, and paleotemperature (see reviews in Clementz, 2012; Koch, 2007).

Knowledge of the ratio of ^13^C to ^12^C (expressed as δ^13^C, the ratio of heavy to light isotope in parts permil (‰) relative to an international standard, here PDB, Pee Dee Belemnite) in bone or tooth hydroxyapatite of herbivores permits estimation of the relative contribution of plants of different photosynthetic groups (C_3_ vs. C_4_/CAM) to an individual’s diet and the reconstruction of its feeding ecology (Bocherens et al., 1996; Cerling et al., 1997, 1998; Fox & Koch, 2004; Franz-Odendaal et al., 2002; Koch, 1998; Koch et al., 2004; Latorre et al., 1997; Macfadden & Cerling, 1996; Zazzo et al., 2000). Additionally, carbon isotope analysis also allows for the identification of taxa or individuals that habitually consumed resources from closed-canopy forests, contributing to our understanding of regional paleoecology (Tejada et al., 2020; Van der Merwe & Medina, 1991). In the analysis presented here, we applied two different diet-hydroxyapatite fractionation values (the isotopic difference between consumer tissues and the foods they consumed), -12‰ and -14.6‰, thus accounting for the range of proposed fractionation values for wild ungulates (Balasse, 2002; Cerling & Harris, 1999; Lee-Thorp & Van der Merwe, 1991; Passey et al., 2005; Sullivan & Krueger, 1981). Generated data were compared with a large (n=1366 representing 79 taxa) modern comparative dataset of Suess-corrected artiodactyl dental enamel δ^13^C values (Wang and Badgley, 2022).

The ratio of ^18^O to ^16^O, or δ^18^O, in biological apatite largely reflects the oxygen isotope composition of water consumed by an organism while alive (Sponheimer & Lee-Thorp, 1999), which, in turn, varies systematically with temperature and precipitation (Dansgaard, 1964; Gat et al., 1996). As such, δ^18^O in animal bioapatite is a widely employed paleothermometer or paleohydrometer allowing for proxy characterization of past climates (Koch, 1998). Observed isotopic values of hydroxyapatite carbonate require several steps of conversion before paleoclimatic determinations can be made (Pryor et al., 2014; Stevens et al., 2011). First, the observed δ^18^O values, which were reported relative to the V-PDB standard, must be converted to the V-SMOW scale, following (Coplen et al., 1983). Next, this converted value is used to derive a value for the oxygen isotope signature of the water consumed/ingested, or δ^18^O_dw_ (Iacumin et al., 1996; Stevens et al., 2011). These values were compared with modern δ^18^O data for the region garnered from the Online Isotopes in Precipitation Calculator (Bowen 2025, Bowen and Revenaugh 2003, Bowen, et al. 2005, IAEA/WMO 2015).

Samples were taken from the apex to cervix of the buccal portion of each crown, thus reflecting an isotopic average of 6–14 months (based on the mineralization times discussed above). Extraction of hydroxyapatite followed protocols first established in Lee-Thorp (1989) and Krueger (1991) and modified by Pestle (2010). Approximately 150 mg (weighed to the nearest 0.1 mg) of ground enamel was placed in a 50 mL centrifuge tube. After weighing, each sample underwent 24 h of organic oxidation using 30 mL of 50% bleach (∼2.5% NaOCl). Samples then were rinsed to neutral pH through a repeated process of centrifugation, decanting, and addition of distilled water. Next, labile carbonates were removed by the addition of 30 mL of 0.1 M acetic acid to each centrifuge tube for a total of four hours, with a 5 min vacuum treatment after 2 h. After the acid treatment, each sample was rinsed to neutral pH before being placed in a 50 °C oven overnight to dry the resulting product. Start and end weights were recorded for all hydroxyapatite samples (weighed to the nearest 0.1 mg) and used to calculate the hydroxyapatite yield (wt%).

Isotopic analysis was performed in the Marine Geology and Geophysics Stable Isotope Laboratory at the Rosenstiel School of Marine and Atmospheric Science, University of Miami. Analysis took place using a Kiel-IV Carbonate Device coupled to a Thermo Finnigan DeltaPlus IRMA, employing an OCC (optically clear calcite) standard calibrated to NBS-19 (δ^13^C=1.9±0.1‰; δ^18^O=-1.6±0.1‰). Standards were analyzed in every sample set at the beginning and end of the run, as well as in-between the analyzed samples to ensure instrumental stability. Check samples of the standard were also run as unknowns in every run to verify measurement accuracy (Szpak et al., 2017). Long-term precision for the analysis of both δ^13^C and δ^18^O was better than 0.1‰. Results for both isotope systems are presented relative to the V-PDB standard (Koch, 2007).

## Statistical Analysis

All visualization and statistical analysis were performed in R v 4.5.1 (R Core Team 2025) using packages *dplyr* (Wickham, et al., 2023) and *ggpubr* (Kassambara 2025). Due to the limited sample size, particularly for the Late Pleistocene, we refrained from conducting inferential statistical tests comparing the two epochs.

## Results

### Bayesian chronological modelling

The modelling identified two major (≥60%) and marginal outliers with posterior probabilities of 64% (Beta-261274) and 42% (Beta-202780), respectively, both located in the mid-to-upper sequence (Figure 2). These values are likely overestimates, potentially resulting from vertical mixing, taphonomic processes, and/or sampling issues. The presence of outliers in this section highlights the need for further research to enhance chronological resolution. The estimated ages for each sample, based on the model, are presented in Table 1.

**Figure 2.**
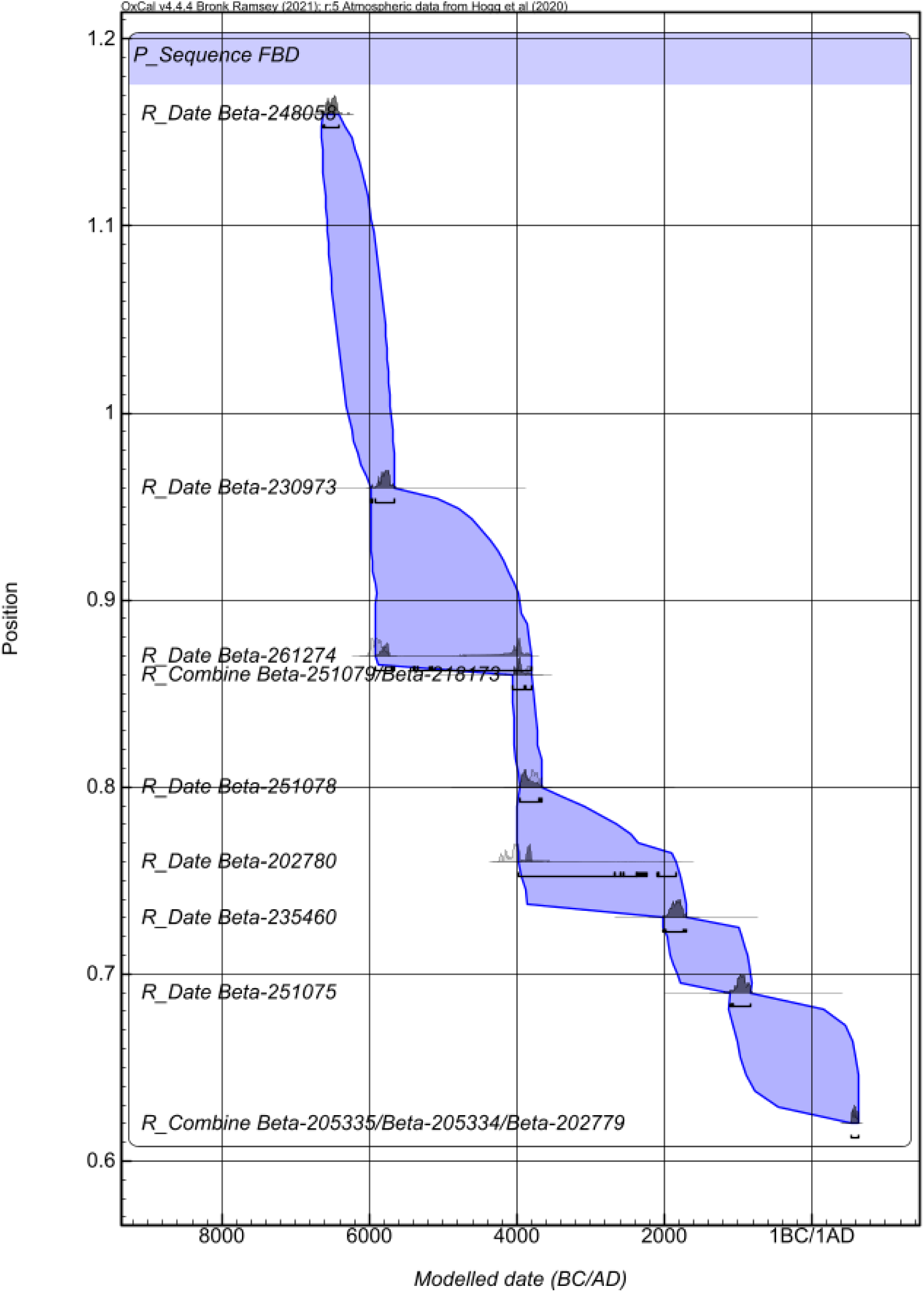
Depositional model for the Holocene facies at Cuvieri Cave. Outlier analysis output is shown next to each laboratory code as ‘[O: posterior probability/prior probability]’. Light grey distributions represent unmodeled, calibrated radiocarbon ages, whereas dark grey distributions show the corresponding modeled age estimates. Black bars underneath each denote the CI at 95.4%, with the plus symbols marking the median. The purple envelope represents the modelled age-depth uncertainty at the 95.4% CI.

### δ^13^C and δ^18^O Isotope Analyses

All samples yielded sufficient material for analysis, with post-treatment hydroxyapatite yields averaging 72.2±6.8%. Observed δ^13^C_PDB_ values ranged from -13.7 to -8.4‰ (mean -10.6±1.3‰), while δ^18^O_PDB_ values averaged -3.3±1.1‰, with an overall range extending from -5.5 to -1.4‰ (Table 1).

Regarding δ^13^C, application of a 12‰ fractionation value yielded an average dietary carbon isotope signature of -22.6±1.3‰, whereas a 14.6‰ fractionation produced an average δ^13^C_diet_ of -25.2±1.3‰. If one considers the latter value, and non-closed canopy plants, the analyzed individuals appear to have consumed almost pure C_3_ plants, which have an average δ13C of 25.5±2‰, with C_4_ plants (e.g., tropical grasses, which average 9.5±1.0‰) forming, on average, only 4.1±4.7% (range 0.0–15.6%) of individual diet. Application of the 12‰ fractionation value yielded higher estimates of C_4_ consumption, averaging 18.2±7.9% (range 0.0–31.9%). If, on the other hand, the δ^13^C of closed-canopy plants (-31‰) is used as the C_3_ endmember, average C_4_ consumption jumps to 27.0±5.9% when using the 14.6‰ fractionation value or 39.1±5.9% when using a 12‰ correction. Regardless of which fractionation value or C_3_ endmember is applied, C_3_ plants formed the majority to preponderance of the analyzed individuals’ diets, with a far smaller contribution coming from C_4_ or CAM plants.

When compared with the enamel δ^13^C of 1366 modern artiodactyls grouped by feeding ecology (Wang and Badgley 2022), the Cuvieri *S. gouazoubira* fall closest to the Browser and Generalist categories, while also overlapping with taxa characterized as Browser-Grazer Intermediate (Figure 3).

**Figure 3.**
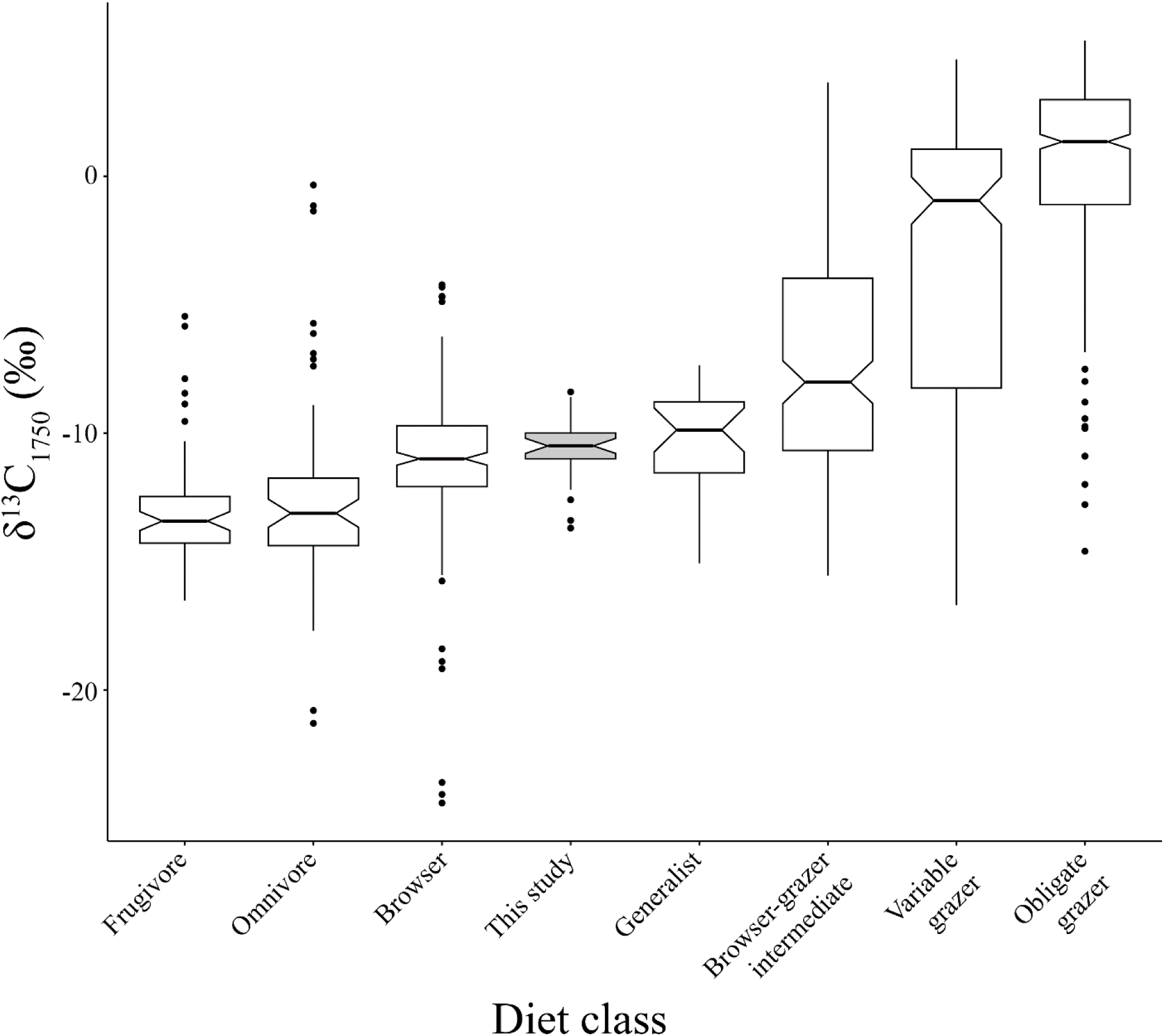
Fossil fuel/Suess corrected carbon isotope values (δ^13^C_1750_) of the dental enamel of Cuvieri *S. gouazoubira* compared with 1366 artiodactyls (Wang and Badgley, 2022), grouped by feeding ecology.

There was no meaningful difference in δ^13^C values between individuals of the Pleistocene and the Holocene (Figure 4). Average Pleistocene δ^13^C was -10.9±1.7‰, while Holocene individuals averaged a very similar -10.5±1.2‰. By extension, there was little difference in the consumption of C_4_ plants between the two epochs. When considered against modeled age, there is no shift in δ^13^C over the time interval represented (Figure 5).

**Figure 4.**
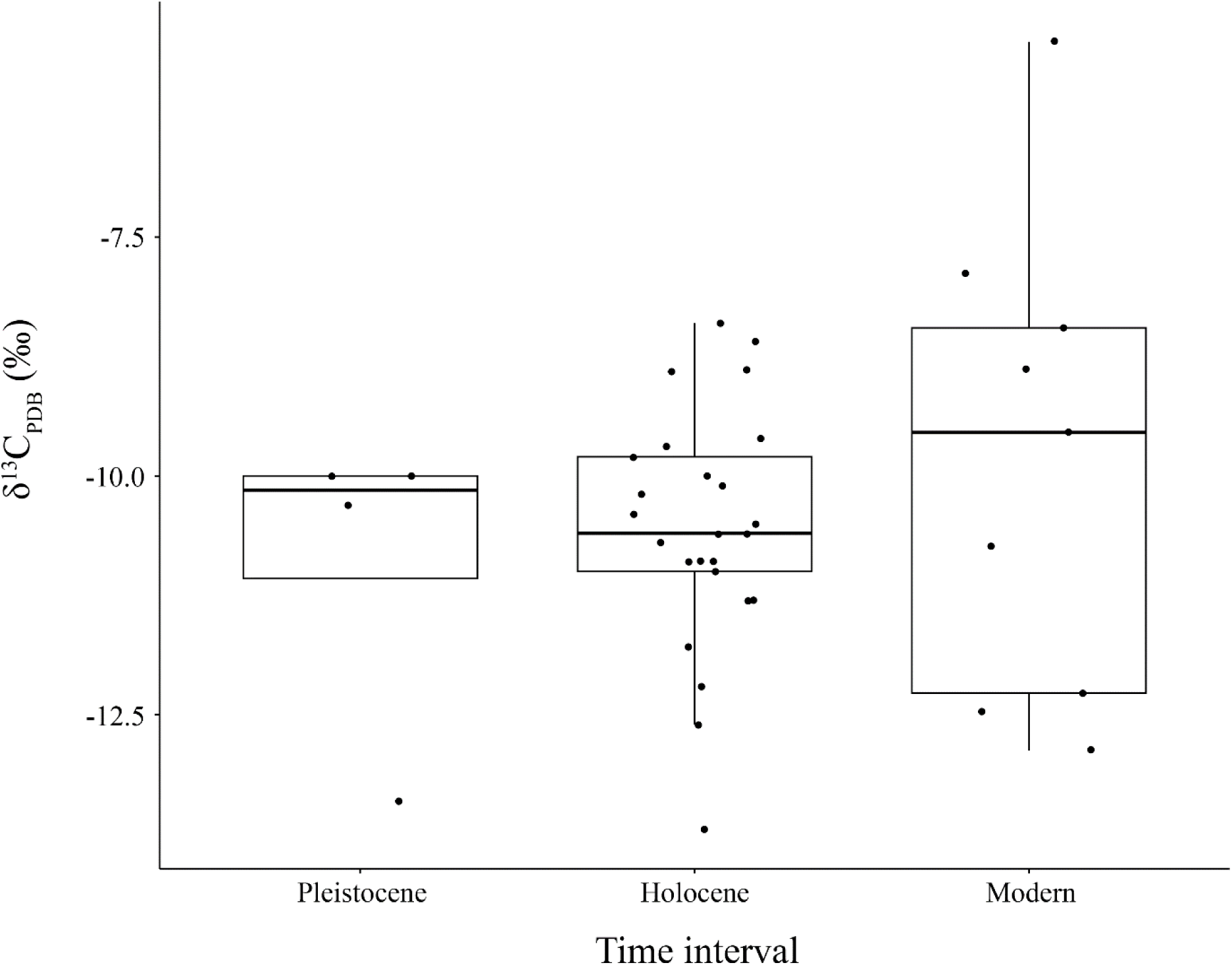
Enamel δ^13^C_PDB_ of Cuvieri and modern *S. gouazoubira*, by time period.

**Figure 5.**
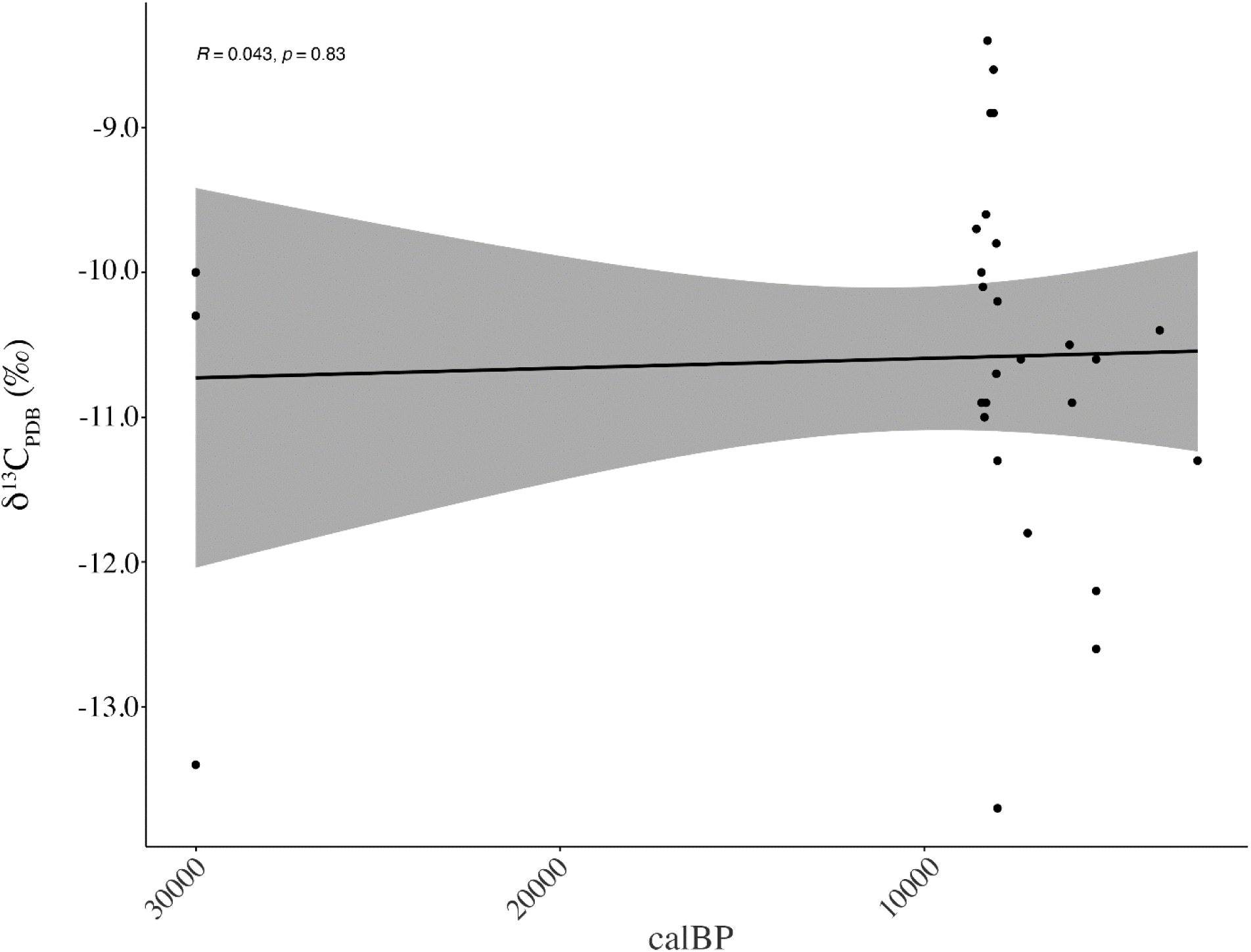
Enamel δ^13^C_PDB_ of Cuvieri *S. gouazoubira* by modeled median age. Pleistocene samples without modeled age were stipulated to be older than 30,000 cal BP following Haddad-Martim et al. (2017).

Both the Pleistocene and Holocene individuals from Cuvieri Cave possess higher mean δ^13^C than nine modern specimens of the taxon (Wang and Badgley 2022: Table S1), although the Cuvieri individuals overlap substantially with the higher portion of the modern range (Figure 4). The modern *S. gouazoubira* with δ^13^C values resembling the Cuvieri specimens (ranging from -5.5 to -10.7‰) come from the Dry Chaco of northwestern Paraguay (approx. 60.1° W 20.8° S), a region of open xerophytic forests, shrublands, and savannah that, based on climate data obtained from geodata/terra has a mean annual temperature of 25.6° C and average rainfall of 630 mm. The isotopic similarity of the Cuvieri individuals with these modern individuals from the Dry Chaco may indicate generalist/browsing feeding behaviors in an open environment, and perhaps one that was drier than current conditions at Cuvieri Cave, which has an average annual temperature of 21.1° C and receives 1330 mm of rain.

Turning to δ^18^O_dw_ (extrapolated, as noted above, from the measured δ^18^O values of apatite carbonate), the overall average was -6.1±1.1‰, with a range that extends from -8.4‰ to -4.2‰. Unlike the case with carbon isotopes, there was a marked difference between the two geological Epochs. The Pleistocene individuals had an average δ^18^O_dw_ of -7.9±0.4‰, while the Holocene individuals averaged -5.8±0.9‰, both of which are lower than the current δ^18^O of precipitation at Cuvieri Cave, which (based on monthly values from OIPC) averages -5.3 ±1.5‰ (Figure 6).

**Figure 6.**
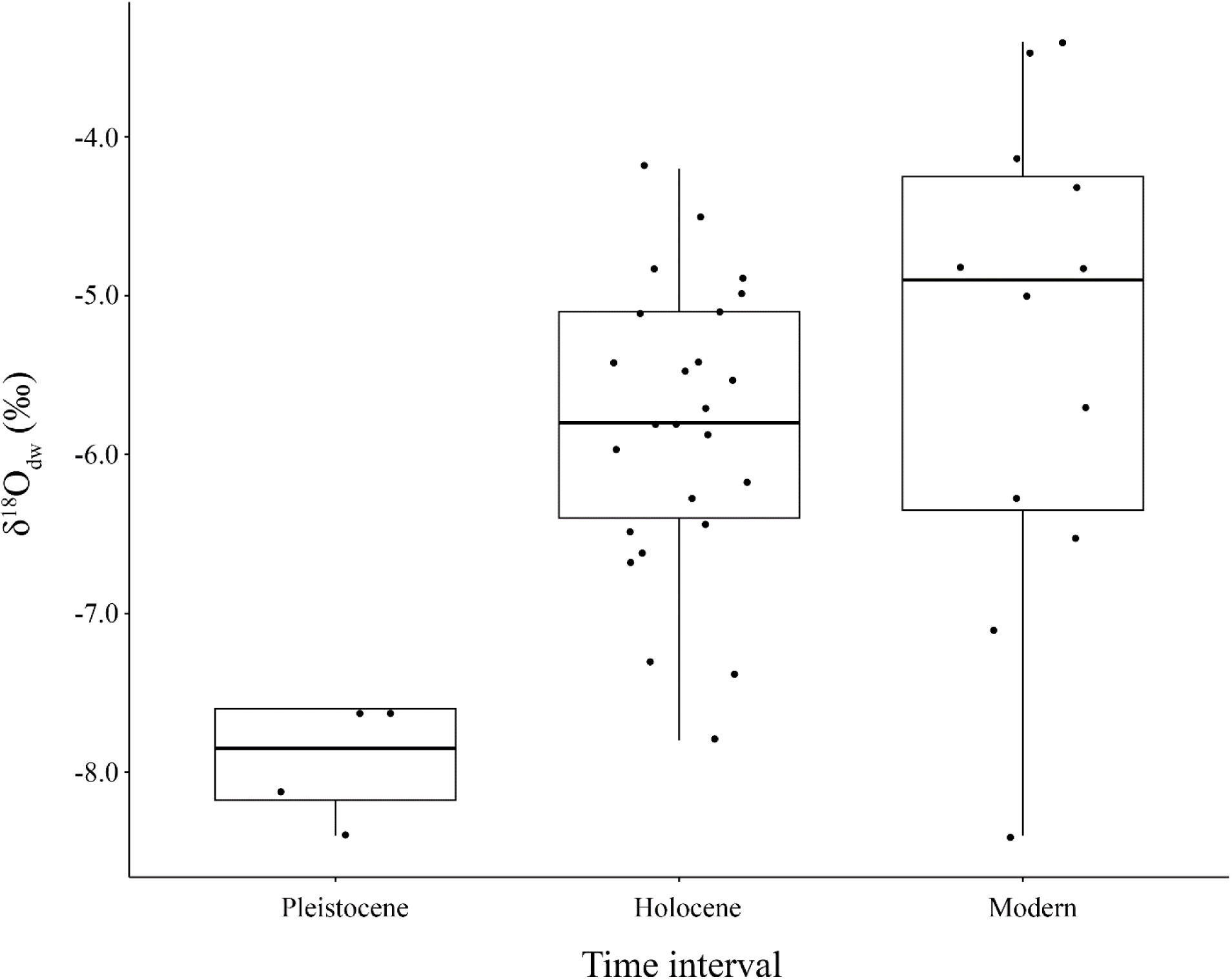
δ^18^O_dw_ derived from Cuvieri *S. gouazoubira* enamel, compared with modern precipitation δ^18^O for region obtained from Online Isotopes in Precipitation Calculator (OIPC).

At lower latitudes (< 30° S/N) there is a complex relationship among temperature, precipitation amount (the amount effect), and oxygen isotopes, such that δ^18^O is not as strictly predictive of temperature as at mid- and high latitudes (Rozanski and Araguás 1995). Indeed, decoupling the effects of changes in temperature and rainfall in the tropics is complex enough that we are reluctant to ascribe the observed shift to anything more specific than a change in prevailing climatic conditions from the late Pleistocene to Holocene to present. The significant diachronic trend towards higher δ^18^O_dw_ is evident also when the brocket deer δ^18^O_dw_ are considered versus modeled age (Figure 7).

**Figure 7.**
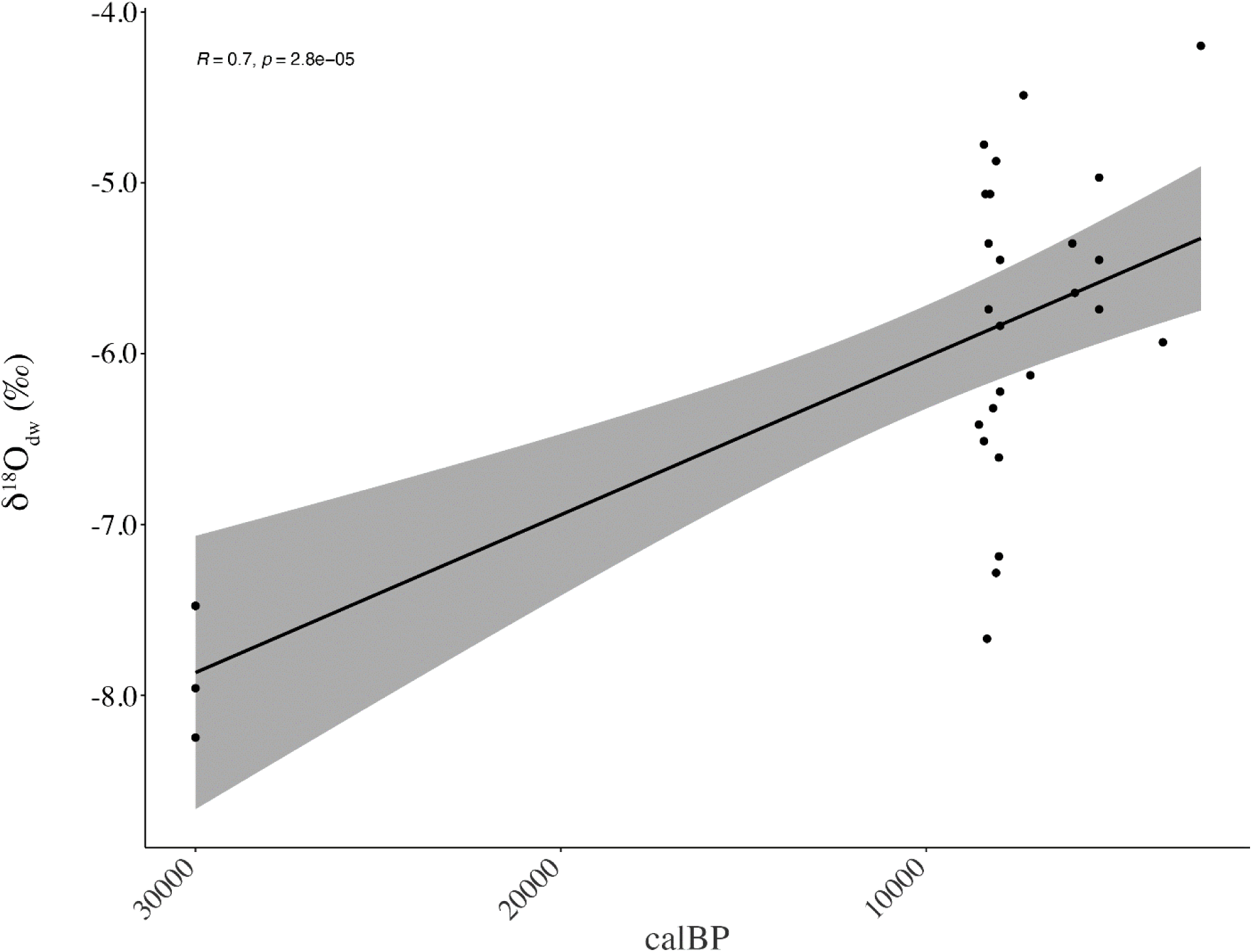
Enamel-derived δ^18^O_dw_ of Cuvieri *S. gouazoubira* by modeled median age. Pleistocene samples without modeled age were stipulated to be older than 30,000 cal BP following Haddad-Martim et al. (2017).

## Discussion

The data presented here, despite their limited sample size, do not indicate a diachronic change in the feeding strategy for the gray brocket deer in the Cerrado between the Late Pleistocene and the Holocene, at least based on the aspects of diet captured in carbon stable isotopes (Figure 4). Our analysis supports gallery/open forest browsing or generalist feeding, perhaps in a more arid environment than seen at present, as the main feeding strategy for all gray brocket deer individuals. This conclusion is consistent with the current understanding of the species’ feeding strategy more broadly (Black-Décima et al., 2010; Black-Décima & Vogliotti, 2016, Grotta-Neto et al., 2024). Our study aligns with the limited research conducted on this taxon in another region of Brazil (Dantas et al., 2020), and with studies of browsers from elsewhere in the southern hemisphere before/after the LGM, which similarly did not observe a significant shift in δ^13^C values between Late Pleistocene and Holocene samples (Sealy et al., 2020). This stands in contrast to studies conducted in the northern hemisphere, which reported differences between epochs (see Sealy et al. [2020] and references therein).

Given the dietary flexibility of gray brocket deer (Black-Décima et al., 2010; Black-Décima & Vogliotti, 2016), and the present and past diversity of plant species in the study region (Raczka et al., 2017), any of the estimates of C_4_ contribution presented above (4.1±4.8% to 39.1±5.9%) seem entirely realistic, as does the variability observed among the Holocene individuals. What is more certain from our results is that, by any fractionation estimate, the plants being consumed by these individuals were not predominantly originating from closed canopy environments, aligning with other isotopic studies (Grotta-Neto et al. 2024) and observational studies of the species’ biology (Black-Décima et al., 2010; Black-Décima & Vogliotti, 2016), but see (Grotta-Neto et al., 2019). Plant tissues from such environments will possess δ^13^C values of -31‰ or lower (Van der Merwe & Medina, 1991), nearly 10‰ lower than the lowest estimated dietary value for the twenty-nine analyzed gray brocket deer individuals (-25.7‰). This lack of evidence for feeding in closed-forest habitats reinforces the interpretation that all sampled individuals indeed represent *S. gouazoubira*, in contrast with the red brocket deer (*Mazama americana*), which is known to inhabit and feed within closed forests (Duarte & Vogliotti, 2016).

Unlike the case of the carbon isotopes, comparison of the Pleistocene and Holocene individuals reveals clear differences in the oxygen isotope composition between epochs (Figure 5). As a consequence, one can make certain, albeit contingent, climatic inferences. Indeed, the complexities of the relationship among temperature, precipitation amount, and oxygen isotopes renders the evidenced trend in climatic change rather equivocal (Rozanski and Araguás 1995).

On the one hand, these data can be seen as consistent with the understanding that the period from the LGM to the mid-Holocene in the Cerrado was cooler and drier than at present (Bissa and Toledo 2015; Salgado-Labouriau, et al., 1997). This interpretation finds support in palynological and geochemical studies of nearby Cerrado locations, which identified a dry and cool climate in the region from around 18,000 – 4000 years ago (Salgado-Labouriau, et al., 1997; Bissa and Toledo 2015) with increases to current levels beginning thereafter.

However, the increase in δ^18^O observed in the transition from the Late Pleistocene to Holocene samples also could be argued to be a consequence of the reduction in precipitation suggested by contemporary stable isotope records observed in speleothems from the Cerrado and other nearby locales (Azevedo et al., 2021; Cruz et al., 2005; Novello et al., 2019). Elsewhere, an increase in herbivore enamel δ^18^O has been argued to be associated with both warmer and drier conditions (Kohn et al., 2022). In this interpretation, the observed 2.1‰ increase in *Subulo* enamel δ^18^O could be associated with a similar magnitude (∼2.5‰) rebound in speleothem δ^18^O seen in speleothem records as the Holocene dry period begins after the conclusion of the Younger Dryas, post 12,900-11,600 years ago (Novello et al., 2017:3).

Thus, multiple reasonable explanations exist for the observed shift in δ^18^O, multiple of which are consistent with different locally available paleoclimatic records. As such, we are most comfortable stating that the shift in δ^18^O seen in the Cuvieri Cave *Subulo* enamel is evidence of a climatic shift, the magnitude and directionality of which is uncertain. What seem irrefutable, however, is that this taxon maintained general dietary stability over this period of changing local climate. In that regard, the data and interpretation presented here provide a useful framework and stimulus for future research on this taxon, as well as on other faunal components of the Cerrado region.

## Supporting information

https://doi.org/10.5281/zenodo.23083803

## Acknowledgements

We are deeply indebted to José Hein for kindly allowing and supporting us during our field seasons on his property, to Maria Mercedes Okumura for providing access to the material housed at the Laboratório de Estudos Evolutivos Humanos, Instituto de Biociências, Universidade de São Paulo. We also thank Leonardo M. Borges for assisting with data from the Cerrado, Max Ernani Cezário and Artur Chahud for their assistance with the material in the Laboratório de Estudos Evolutivos Humanos, and IBAMA, ANM, and IPHAN for providing the permits necessary for our investigations. Finally, we thank Romain Amiot and an anonymous reviewer for providing constructive feedback that substantially improved our work.

## Data availability

Supplementary Information 1 is available in Zenodo (https://doi.org/10.5281/zenodo.23083803)

## Authors contributions

WJP: Conceptualization, Formal analysis, Investigation, Data Curation, Funding acquisition, Supervision, Writing - Original Draft, Writing - Review & Editing; AH: Conceptualization, Investigation, Funding acquisition, Writing - Original Draft, Writing - Review & Editing; MH: Conceptualization, Writing - Review & Editing; AA: Investigation, Writing - Review & Editing; ENC: Investigation, Writing - Review & Editing; REO: Investigation, Writing - Review & Editing; EDPS: Investigation, Writing - Review & Editing; LBV: Investigation, Writing - Review & Editing; JMBD: Investigation, Resources, Writing - Review & Editing; WAN: Funding acquisition, Writing - Review & Editing.

