## Supplementary material for "Late Quaternary millennial-scale stability of the feeding strategy of the gray brocket deer [*Subulo gouazoubira* (G. Fischer, 1814); Mammalia] in Southeastern Brazil": https://doi.org/10.5281/zenodo.23083803

Supplementary Information

OxCal code

Plot()

{

Curve("SHCal20","shcal20.14c");

Outlier_Model("General",T(5),U(0,4),"t");

P_Sequence("FBD",100,100,U(-2,2))

{

Boundary();

R_Date("Beta-248058",7690,50)

{

Outlier(0.05);

z=1.16;

};

R_Date("Beta-230973",6930,40)

{

Outlier(0.05);

z=0.96;

};

R_Date("Beta-261274",7050,50)

{

Outlier(0.05);

z=0.87;

};

R_Combine("Beta-251079/Beta-218173")

{

R_Date("Beta-251079",5150,50);

R_Date("Beta-218173",5200,50);

Outlier(0.05);

z=0.86;

};

R_Date("Beta-251078",5050,40)

{

Outlier(0.05);

z=0.8;

};

R_Date("Beta-202780",5250,50)

{

Outlier(0.05);

z=0.76;

};

R_Date("Beta-235460",3550,40)

{

Outlier(0.05);

z=0.73;

};

R_Date("Beta-251075",2830,40)

{

Outlier(0.05);

z=0.69;

};

R_Combine("Beta-205335/Beta-205334/Beta-202779")

{

R_Date("Beta-205335",220,40);

R_Date("Beta-205334",2050,40);

R_Date("Beta-202779",1960,40);

Outlier(0.05);

z=0.62;

};

Boundary();

};

};
